# A yeast model reveals the SMAD3 nuclear import mechanism

**DOI:** 10.1101/2025.07.09.663975

**Authors:** Delfina P. González, Morgan C. Emokpae, C. Patrick Lusk, Mustafa K. Khokha

**Affiliations:** Department of Pediatrics and Department of Genetics, Yale School of Medicine; Department of Cell Biology, Yale School of Medicine; Pediatric Genomic Discovery Program

**Keywords:** TGF-β, SMAD, nuclear transport, importin-7/8, importin-β, nuclear localization signal

## Abstract

TGF-β signaling is important for patterning tissues during embryonic development and relies on the nuclear import of SMADs. Basal SMAD nuclear import mechanisms remain to be fully understood and could benefit from advances in our understanding of nuclear transport machinery. To assess the molecular determinants of SMAD nuclear transport, we take advantage of the unicellular *S. cerevisiae* model that lacks the TGF-β signaling complexity of vertebrates but has conserved nuclear transport machinery. In this minimal system, we find that the steady-state distribution of SMAD2/3, SMAD4, and SMAD1/5 are Ran-dependent and either accumulate in, or are depleted from the nucleus. Conditionally inhibiting each of the karyopherin/importins demonstrates that SMAD3 is imported by orthologues of importin-β, importin-7 and importin-8. The previously defined nuclear localization signal is insufficient to confer nuclear import. Instead, our data suggest the entire MH1 domain may act as an importin binding surface. Deleting a portion of this domain impairs SMAD3 function in *Xenopus* and SMAD3 nuclear enrichment in HEK293T cells. Thus, the yeast platform provides an efficient strategy to illuminate the nuclear transport mechanisms of embryonic signaling effectors.

## INTRODUCTION

TGF-β signaling is critical for embryonic patterning. Binding of a TGF-β family ligand to the Type II receptor stimulates dimerization with a Type I receptor. Dimerization causes phosphorylation of Receptor SMADs (R-SMADs) and subsequent complex formation with the co-SMAD. Current models suggest that this phosphorylation and complex formation leads to enrichment of R-SMADs in the nucleus and activation of target genes^[1],[2],[3]^. Mis-regulation of these signaling pathways can lead to multiple diseases including birth defects and cancer^[4]^. Therefore, a better understanding of the TGF-β signaling pathway could maximize its therapeutic potential, in particular the nuclear transport of SMADs.

In general, nuclear transport of proteins requires the recognition of unstructured signal sequences, either nuclear localization or nuclear export signals (NLSs and NESs) by one of the ~20 nuclear transport receptors (NTRs) of the β-karyopherin family (also called importins and exportins). NTRs ferry cargo through the transport channel of the nuclear pore complex (NPC) with directionality and energy imparted by the Ran GTPase. Ran is found predominantly in a GTP-bound state in the nucleus and as Ran-GDP in the cytosol. The binding of Ran-GTP to import or export cargo complexes leads to their dissociation or stabilization, respectively^[5]^.

The foundational studies of SMAD nuclear transport preceded the discovery of the complete repertoire of NTRs that we have now. One major advance was the discovery in *D. melanogaster* that the Importin-7/8 ortholog, moleskin (Msk), impacts SMAD1, 2, and 3 nuclear enrichment. This interaction was identified by an RNAi screen for players in Drosophila MAD nuclear import and was confirmed by genetic and biochemical methods for both Msk and human importin-7 and 8^[6]^. Some earlier studies identified a role for importin-β specifically in activated SMAD3 nuclear enrichment^[7],[8]^. However, whether other more recently described NTRs also play a role in SMAD nuclear transport remains an open question.

With regard to the SMAD NLS, early studies identified a classical “c” NLS-like sequence “KKLK” (modeled off of the SV-40 T-Antigen NLS) that might be recognized by the Importin-α/β heterodimer^[9],[10],[11]^. Close inspection of these data, however, indicate that although this sequence was necessary for SMAD import, it was insufficient to confer nuclear localization of a reporter. Thus, it failed to meet the formal criteria for an NLS suggesting that other sequences or structural elements may also be playing a role in SMAD nuclear import^[10],[12],[13]^. Consistent with this idea, mutating the KKLK sequence to AALA did not impact SMAD’s ability to interact with Importin-7 or 8^[13]^. Thus, a comprehensive screen of all known NTRs and an unbiased reexamination of any potential SMAD NLS sequences may provide additional insight into SMAD nuclear transport mechanisms.

One challenge in understanding SMAD nuclear transport is its dynamic nature in animal cells and severe alterations in cancer cell lines. Therefore, we sought to introduce a minimal system, which allows us to focus solely on nuclear transport without the complexities of the vertebrate system. The yeast *Saccharomyces cerevisiae* shares a conserved nuclear transport system with humans **(Table 1)** but lacks the TGF-β/SMAD signaling pathway and other multicellular signaling complexities that can confound conclusions about the nuclear transport of a single signaling component. Therefore, by assessing SMAD localization in yeast, we can focus on the mechanism of nuclear transport in a mimic of the basal condition by leveraging the wealth of genetic tools available for this system.

**Table 1.** Yeast Orthologs of Human Nuclear Transport Receptors.

| Yeast Ortholog | Human NTR |
| --- | --- |
| Kap95 | Kap-β1 |
| Kap104 | Transportin1 |
| Kap108/Sxm1 | Importin-8/RanBP8 |
| Kap111/Mtr10 | Transportin3-SR/TNPO3 |
| Kap114 | Importin-9 |
| Kap119/Nmd5 | Importin-7/RanBP7 |
| Kap120 | Importin-11 |
| Kap121 | Importin-5/ Kap-β3 |
| Kap122/Pdr6 | Importin-13 |
| Kap123 | Importin-4/RanBP5 |
| Kap60/Srp1* | Kap-α |
| Xpo1* | CRM1/Exportin-1 |
| Los1* | Exportin-t |
| Kap142/Msn5* | Exportin-5 |
| Cse1* | CAS |

\*Not part of the anchor away screen.
Adapted from [5]

We found that at steady-state eGFP-tagged nodal/activin SMADs (SMAD2/3) enrich in the yeast nucleus while bone morphogenetic protein (BMP) SMADs (SMAD1/5) and the Co-SMAD (SMAD4) are depleted in a Ran-cycle dependent manner. We then focused on SMAD3, in which our unbiased screen revealed that Kap95 (importin-β), Kap108 (importin-8) and Kap119 (importin-7) are required for its nuclear accumulation. Finally, we determined that the entire MH1 domain is the minimally sufficient sequence necessary for optimal SMAD3 nuclear enrichment. In our yeast model, a construct containing the previously identified KKLK sequence localizes to the cell periphery suggesting that it is not the SMAD3 NLS. To test requirement, we deleted 20 amino acids in the MH1 domain (which does not include the KKLK sequence) which disrupts SMAD3 nuclear enrichment in HEK293T nuclei and is required for SMAD signaling in *Xenopus*. These results suggest that basal SMAD3 nuclear import relies on an interaction with the yeast orthologues of importin-β, importin-7, and importin-8 through a non-conventional NLS within a structured domain. Our results demonstrate the utility of yeast for isolating and interrogating the nuclear transport step of developmental signaling effectors that could be broadly applicable for any vertebrate transcription factor.

## RESULTS

### SMAD2/3 (Nodal/Activin), SMAD1/5 (BMP), and SMAD4 localize differently at steady state

We first tested how N-terminally eGFP-tagged human SMADs would distribute in *S. cerevisiae*. We integrated an eGFP, eGFP-cNLS, or eGFP-SMAD sequence into the yeast genome at the *URA3* locus under the control of a constitutive yeast *ADH1* promoter. A cNLS-GFP reporter acts as a positive control as its energy-dependent import mechanism is known to be mediated by Importin-α/Importin β1^[5]^. We transformed these constructs into a wild-type yeast strain expressing Heh2-mCherry, an inner nuclear membrane protein that serves as a nuclear envelope marker^[14]^.

eGFP was evenly distributed between the nucleus and the cytoplasm, while eGFP-cNLS was enriched in the nucleus (**Fig. 1A**). eGFP-SMAD2/3 was enriched in the nucleus, while eGFP-SMAD4 and eGFP-SMAD1/5 were depleted. eGFP-SMAD9 was evenly distributed between the nucleus and cytoplasm (**Fig. 1A**). We calculated a nuclear to cytoplasmic (N/C) ratio from nuclear and cytoplasmic fluorescence intensity measurements of each strain (**Fig. 1B**). These results indicated that the Nodal/Activin SMADs are relatively enriched in the nucleus while the BMP SMADs and Co-SMAD were depleted. These results are consistent with current models of SMAD localization with regards to co-SMAD4, which is depleted from the nucleus in the absence of signaling, but not for the R-SMADs^[6],[15],[16]^. Importantly, these SMAD localizations are under basal conditions.

**Figure 1.**
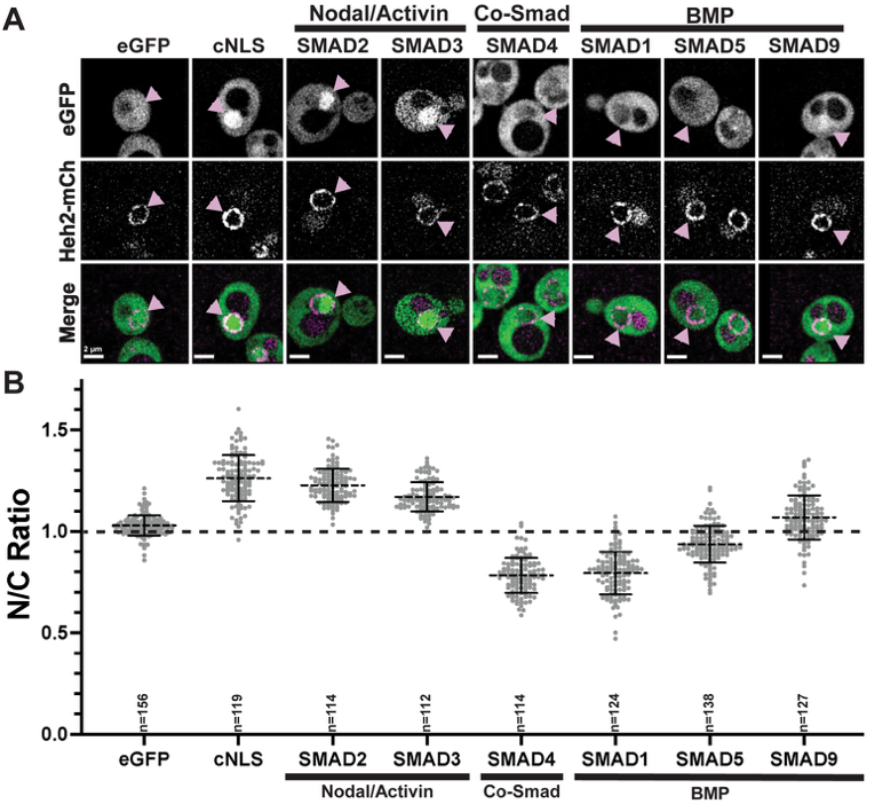
SMAD2/3 (Nodal/Activin), SMAD1/5 (BMP), and SMAD4 have different nuclear enrichments. **(A)** Images of WT yeast co-expressing the nuclear envelope protein Heh2-mCherry and either eGFP, eGFP-cNLS, and eGFP-SMAD. Pink arrows mark the same location across merged and single channel images. Scale bar= 2 μm. **B)** Individual nuclear-to-cytoplasmic ratios of eGFP, eGFP-cNLS, and eGFP-SMAD constructs expressed in WT yeast. Solid error bars represent standard deviation. Dashed line through scatterplot represents the mean (eGFP 1.0±0.05, eGFP-cNLS 1.3±0.12, eGFP-SMAD2 1.2±0.08, eGFP-SMAD3 1.2±0.07, eGFP-SMAD4 0.8±0.09, eGFP-SMAD1 0.8±0.11, eGFP-SMAD5 0.9±0.07, eGFP-SMAD9 1.1±0.11). Experiments were done in triplicate and the total number of nuclei are labeled in the graph. Dashed line across the graph marks N/C ratio of 1.

### SMAD2/3, SMAD1/5, and SMAD4 nuclear enrichments or depletions are Ran-dependent

To test if the steady state distributions of the SMADs required the Ran system, we leveraged the *mtr1-1* allele, which encodes for a temperature-sensitive Ran-GEF (*SRM1/PRP20/MTR1*)^[17]^. Incubating this strain at the restrictive temperature (37ºC) disrupts the Ran gradient and thus active nuclear transport. At the permissive temperature (RT, 22-25ºC), N/C ratios of all constructs mirrored those in WT cells (**Fig. 2 A, C**, blue). We chose not to test SMAD9 because it did not have a strong nuclear enrichment or depletion. At 37ºC, eGFP stayed evenly distributed. eGFP-cNLS, eGFP-SMAD2/3, and eGFP-SMAD1/5 became more evenly distributed between the nucleus and cytoplasm. eGFP-SMAD4 went from being depleted from the nucleus to being slightly enriched (**Fig. 2B, C**, red). These results indicate that nuclear enrichment of Nodal/Activin SMADs and depletion of co-SMAD4 and BMP SMADs are all Ran-dependent.

**Figure 2.**
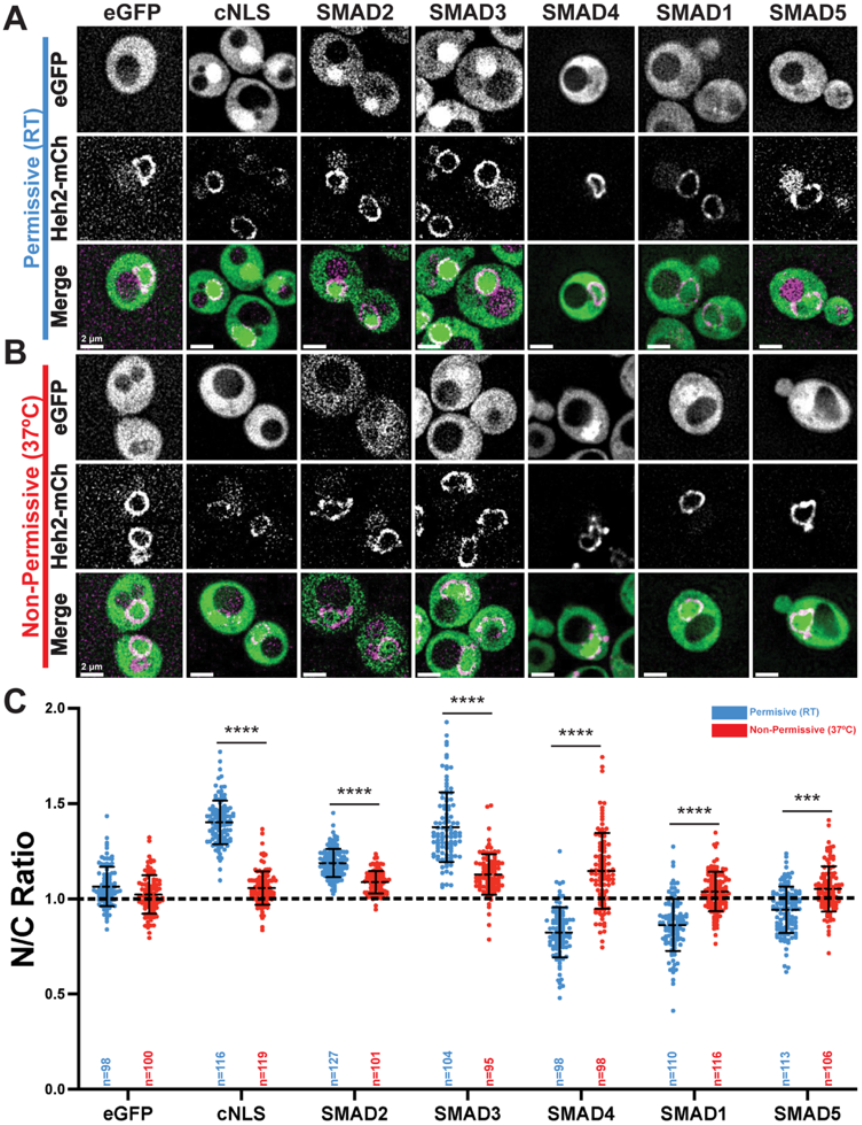
Ran cycle sensitivity of SMAD2/3, SMAD1/5, and SMAD4 nuclear localizations. **(A-B)** Images of the Ran-GEF temperature-sensitive mutant yeast strain (*mtr1-1*) co-expressing the nuclear envelope protein Heh2-mCherry and either eGFP, eGFP-cNLS, and eGFP-SMAD. Scale bar=2 μm. **(A)** Cells imaged at the permissive temperature (Room Temperature). **(B)** Cells imaged after the 1 hr incubation at the non-permissive temperature (37ºC). **(C)** Individual nuclear-to-cytoplasmic ratios of eGFP-tagged constructs expressed in *mtr1-1* yeast. Dots in blue represent N/C ratios of single cells incubated at RT. Dots in red represent N/C ratios of single cells imaged following the 1hr incubation at 37ºC. **** p<0.0001, *** p<0.0002 comparing between RT and 37ºC treatment for each strain (Unpaired T-test). Solid error bars represent standard deviation. Dashed line through scatterplot represents the mean (RT/37ºC eGFP 1.1±0.10/1.0±0.10, eGFP-cNLS 1.4±0.14/1.1±0.09, eGFP-SMAD2 1.2±0.07/1.1±0.06, eGFP-SMAD3 1.4±0.18/1.1±0.11, eGFP-SMAD4 0.8±0.13/1.1±0.20, eGFP-SMAD1 0.9±0.14/1.0±0.10, eGFP-SMAD5 0.9±0.12/1.1±0.12). Experiments were done in triplicate and the total number of nuclei are labeled in the graph. Dashed line across the graph marks N/C ratio of 1.

Of note, loss of the Ran gradient shifted co-SMAD4 from being depleted from the nucleus to being enriched, a result which we currently cannot explain but may reflect an affinity for a nuclear factor. Overall, these results suggest that nuclear enrichment or depletion of these signaling effectors is Ran-dependent and therefore likely relies on an NTR. Additionally, these mechanisms are different between the classes of SMADs. Therefore, we decided to re-visit the nuclear transport mechanism of one of these SMADs, SMAD3, in more depth using unbiased screens to identify the NTR responsible for its import and the NLS it uses to interact with this NTR.

### SMAD3 nuclear enrichment is Kap95, Kap119, and Kap108 (importin-β, importin-7 and importin-8)-dependent

We decided to focus on SMAD3 based on its importance in embryonic development and our interest in left-right patterning in which Nodal signaling plays an important role^[18], [19], [20], [21], [22], [23]^. To deplete NTRs, we used the anchor away approach^[24]^. This approach harnesses the rapamycin-induced formation of a complex between FKBP12 and FRB. This inducible system can rapidly sequester essential cytoplasmic and nuclear proteins. We designed our inducible system to rapidly sequester NTR-FRBs to the plasma membrane using the abundantly expressed Pma1-FKBP12. This approach has previously been used to identify the NTR for β-catenin in WNT signaling^[25]^.

In a negative control strain that lacks an FRB-tagged NTR and all the other tested FRB-tagged NTR strains, treatment with rapamycin had no effect on eGFP-SMAD3 nuclear enrichment (**Fig. 3B, Fig. S1**). Meanwhile, membrane trapping of Kap95-FRB, Kap108-FRB, and Kap119-FRB altered eGFP-SMAD3 nuclear enrichment (**Fig. 3A, B**), consistent with previous studies^[6],[7],[8],[13]^. This validates our yeast system for studying the import of vertebrate signaling effectors. We next used a deletion analysis to identify the sequence elements that contribute to SMAD3’s nuclear accumulation.

**Figure 3.**
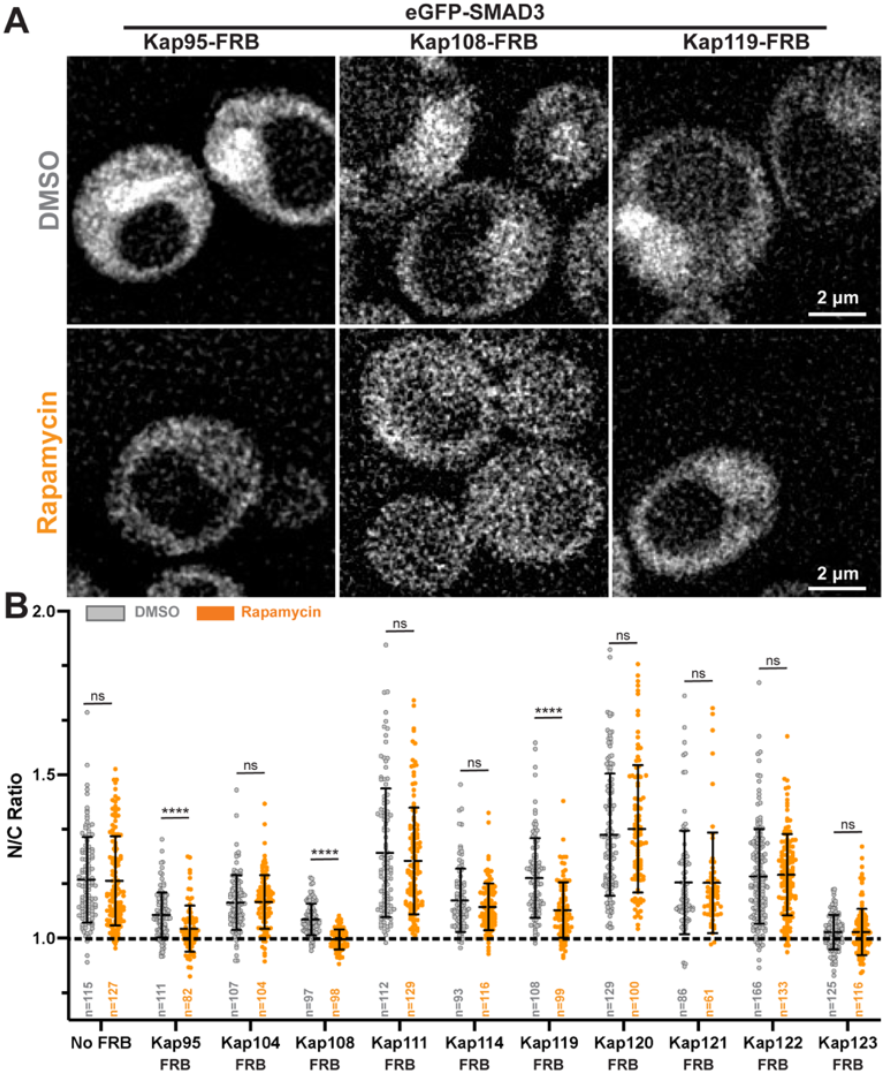
SMAD3 nuclear enrichment depends on Kap95, Kap108, and Kap119. **(A-B)** Images of select anchor-away yeast strains co-expressing the nuclear envelope protein Heh2-mCherry and eGFP-SMAD3. The top rows represent cells imaged following a 15 min incubation with DMSO (gray). The bottom rows represent cells imaged following a 15 min incubation with Rapamycin (orange). Scale bar= 2μm. **(A)** Depletion of Kap95, Kap108, Kap119 reduces eGFP-SMAD3 nuclear enrichment. See Supplementary Fig S1 for remaining experimental images. **(B)** Individual nuclear-to-cytoplasmic ratios of eGFP-SMAD3 expressed in anchor-away yeast strains. For a negative control, eGFP-SMAD3 was tested with no FRB-tagged Kaps (No FRB). Dots in gray represent N/C ratios of single cells incubated in DMSO. Dots in orange represent N/C ratios of single cells incubated in rapamycin. **** p<0.0001, *** p<0.0002, ** p<0.002, *p<0.02 comparing between DMSO and Rapamycin treatment for each strain (Unpaired T-test). Solid error bars represent standard deviation. Dashed line through scatterplot represents the mean (NoFRB-SMAD3 1.2±0.13/1.2±0.14, Kap95-SMAD3 1.1±0.07/1.0±0.07, Kap104-SMAD3 1.1±0.08/1.0±0.08, Kap108-SMAD3 1.1±0.05/1.0±0.03, Kap111-SMAD3 1.3±0.20/1.2±0.16, Kap114-SMAD3 1.1±0.10/1.1±0.07, Kap119-SMAD3 1.2±0.12/1.1±0.09, Kap120-SMAD3 1.3±0.19/1.3±0.19, Kap121-SMAD3 1.2±0.16/1.2±0.15, Kap122-SMAD3 1.2±0.14/1.2±0.12, Kap123-SMAD3 1.0±0.05/1.0±0.07). Experiments were done in triplicate. Total number of nuclei are labeled in the graph. Dashed line across the graph marks N/C ratio of 1.

### The SMAD3 nuclear localization signal is not a classical NLS

To identify the sequence elements in SMAD3 that confer nuclear localization, we made several eGFP-SMAD3 constructs guided by previously reported SMAD crystal structures of the MH1 and MH2 domains at locations least likely to disrupt secondary structure (**Fig. 4A**) ^[26–32]^. These constructs were then transformed into WT yeast co-expressing Heh2-mCh, as in Figure 1. The MH1 (1-136) domain had the strongest nuclear enrichment of the 3 domains (**Fig. 4A’, 4D)**. This suggested that the main element driving SMAD3 into the nucleus resides in the MH1 domain, consistent with previous studies of various SMADs^[10],[11],[12]^.

**Figure 4.**
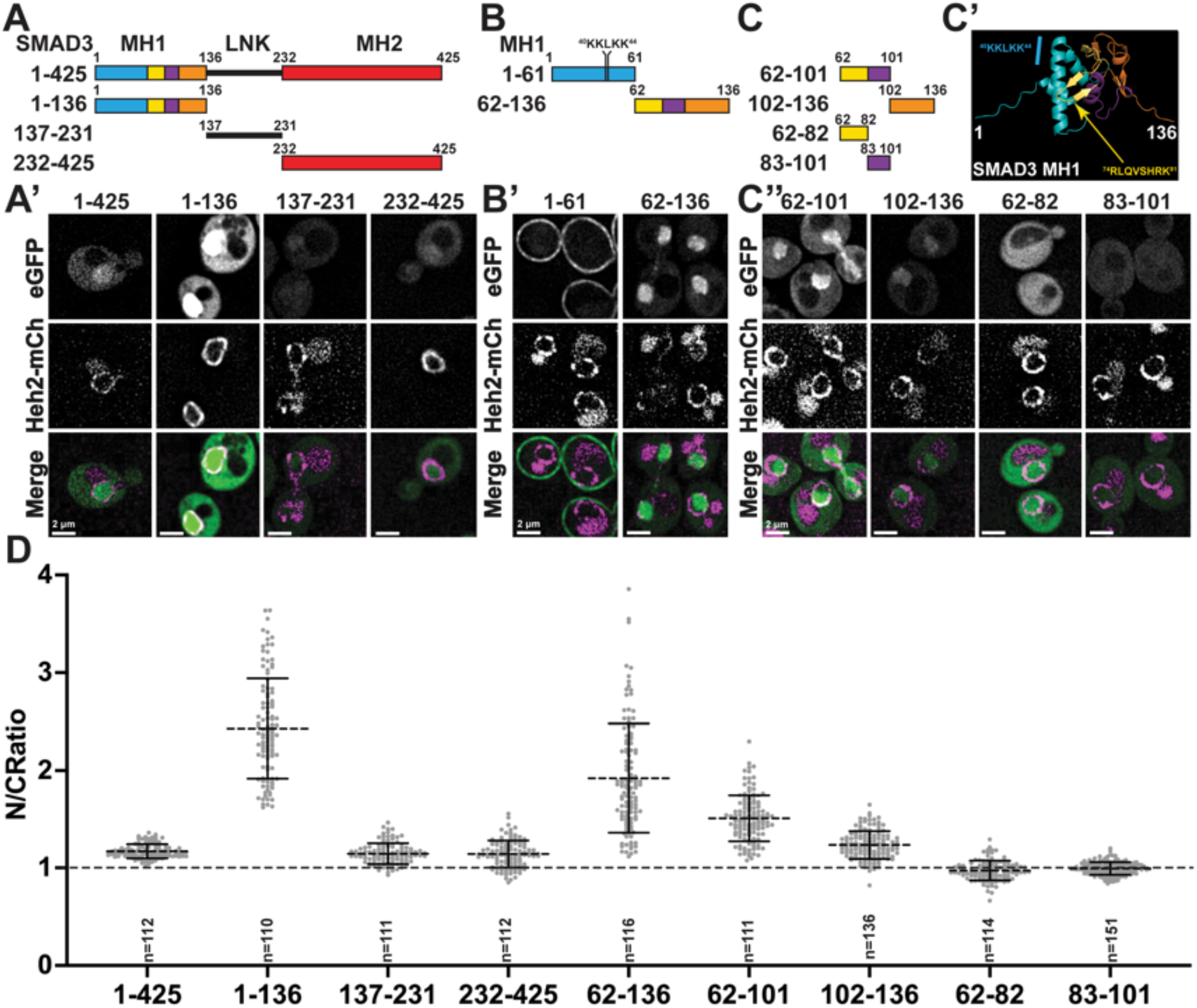
The SMAD3 NLS resides in the MH1 domain. (A)SMAD major domains. The full length human SMAD3 has 425 amino acids and includes an MH1 domain (1-136, turquoise, yellow, purple, orange), a linker domain (137-231, black line), and an MH2 Domain (232-425, red). All constructs are N-terminally tagged with eGFP. **(A’)** Images of WT yeast co-expressing the nuclear envelope protein Heh2-mCherry and one of the eGFP-SMAD3 major domain constructs. Scale bar= 2μm. (B)SMAD3 MH1 N- and C-terminal domains. The human SMAD3 MH1 domain (1-136) was sub-divided between an N-terminal half (1-61, turquoise) and a C-terminal half (62-136, yellow, purple, orange). Not to scale. **(B’)** Images of SMAD3 MH1 N- and C-terminal domains. Images of WT yeast co-expressing the nuclear envelope protein Heh2-mCherry and either the eGFP-SMAD3 N- and C-terminal domain. Scale bar= 2μm. (C)SMAD3 MH1 C-terminal subdomains. SMAD3 (62-82, yellow) contains the β-sheet containing key DNA-binding residues, (83-101, purple) is an α-helix, and 102-136 contains the remainder of the MH1 domain (orange). Not to scale. **(C’)** AlphaFold predicted structure of the SMAD3 MH1 domain. Colors in structure correlate with schematics in A, C, and D. Structure is annotated with features described in B and C. The N-terminus is on the left (1) and the C-terminus in on the right (136)[46],[47]. **(C’’)** Images of SMAD3 MH1 c-terminal subdomains. Images of WT yeast co-expressing the nuclear envelope protein Heh2-mCherry and one of the eGFP-SMAD3 constructs. Scale bar= 2μm. (D)Nuclear-to-cytoplasmic ratios of all eGFP-SMAD3 constructs expressed in WT yeast. Solid error bars represent standard deviation. Dashed line through scatterplot represents the mean (1-425 Full 1.2±0.07, 1-136 MH1 2.43±0.51, 137-231 Linker 1.15±0.11, 232-245 MH2 1.14±0.14, 62-136 1.92±0.56, 62-101 1.51±0.24, 62-82 0.97±0.10, 83-101 0.99±0.07). Experiments were done in triplicate. The total number of nuclei are labeled in the graph. Dashed line across the graph marks N/C ratio of 1. Images and quantification for SMAD3(1-425) are from Figure 1 and presented here for comparison.

Typically, NLSs are short sequences in intrinsically disordered regions (IDR) that bind to an NTR^[5]^. As the SMAD3 MH1 domain is well structured with few IDRs, we were somewhat surprised that it enriched in the nucleus, suggesting that its nuclear localization elements might comprise regions with secondary or tertiary structure. Therefore, we further subdivided the globular MH1 domain at loops connecting secondary structure elements SMAD3 (1-61) includes an unstructured region as well as 2 α-helices, one of which contains the putative cNLS (^40^KKLKK^44^) **(Fig. 4B, C’)**^[10]^. SMAD3 (62-136) includes a β-sheet that contains the DNA binding domain (^74^RLQVSHRK^81^), an α-helix, and a relatively unstructured domain at the N-terminus **(Fig. 4C’)**. eGFP-SMAD3 (1-61) localizes almost exclusively to the cell periphery (**Fig. 4B’**). In contrast, eGFP-SMAD3 (62-136) enriches in the nucleus (**Fig. 4B’, D**). Thus, the region of SMAD3 encompassing 62-136 is sufficient to confer nuclear accumulation but to a lesser extent than the entire MH1 domain suggesting that this sequence does not comprise the entire nuclear localization determinants. Additionally, the previously described NLS containing the^40^KKLKK^44^ does not drive eGFP to the nucleus in our assay.

### The entire MH1 domain is required for optimal nuclear localization

To further refine the SMAD3 NLS, we subdivided SMAD3(62-136) into 2 sub-fragments using SMAD3 crystal structures as a guide (**Fig. 4C, C’**) (PDB: 5od6, 5odg, 6zmn, 1ozj,1mhd)^[27],[28],[31],[32]^. When we divide eGFP-SMAD3(62-136), neither eGFP-SMAD3(62-101), which includes a β-sheet and an α-helix, nor eGFP-SMAD3(102-136), which is relatively unstructured, enrich in the nucleus to the same degree as SMAD3(62-136) although eGFP-SMAD3(62-101) has more enrichment relative to eGFP-SMAD3(102-136) **(Fig. 4C”, D)**. We further subdivided SMAD3(62-101) into fragments containing the β-sheet (62-82) and the α-helix (83-101). These two constructs did not enrich eGFP in the nucleus **(Fig. 4C-D)**. These results suggest that the β-sheet (62-82) and the α-helix (83-101) are a minimal sequence sufficient to drive eGFP into the nucleus. Summarizing our results for putative NLS mapping, there may be multiple sequences in the MH1 domain required for robust nuclear import and some sequences are also sufficient. However, loss of these sequences leads to progressive loss of nuclear enrichment. Therefore, we conclude that although there are sequences that are sufficient to confer nuclear import and may act as NLSs, the entire MH1 is required for efficient nuclear import.

### The SMAD3 (83-101) region is required for nuclear localization in vertebrate models

We next tested the functional necessity of the identified MH1 NLS in the vertebrate developmental model *Xenopus tropicalis*. Since this region contains the DNA binding element, we deleted a portion of the MH1 NLS which did not contain either the KKLKK or the DNA-binding element. We then tested this construct in functional assays. Overactivation of Nodal signaling disrupts left-right patterning leading to changes in cardiac looping and has been well-studied in *Xenopus*^[21],[35]^. We injected 50 pg of either eGFP, eGFP-SMAD3, or eGFP-SMAD3Δ83-101 mRNA with 100-250pg of an mCherry mRNA tracer into a one-cell *X. tropicalis* embryo (**Fig. 5A, B**). Overexpression of full-length eGFP-SMAD3 led to abnormal heart looping in about 22% of embryos compare to just 2% in control eGFP injected embryos. Injection of eGFP-SMAD3Δ83-101 mRNA led to abnormal heart looping in only 6% of embryos **(Fig. 5C)**. In order to determine if this reduction in abnormal cardiac looping is due to a loss of nuclear localization versus a loss in protein function, we added the cNLS (PKKRKV) to the N-terminus of eGFP-SMAD3Δ83-101 **(Fig. 5A)**. Injection of eGFP-SMAD3Δ83-101+cNLS led to abnormal heart looping in 17% of embryos indicating that the cNLS rescues signaling **(Fig. 5C)**. Thus, we conclude that the 83-101 deletion alters nuclear enrichment rather than other protein functions. We next tested these constructs in mammalian cells for nuclear accumulation. eGFP was evenly distributed between the nucleus and cytoplasm and eGFP-SMAD3 enriched in the nuclei (**Fig. 5D**). eGFP-SMAD3Δ83-101 led to a loss of nuclear enrichment, but eGFP-SMAD3Δ83-101+cNLS regained enrichment as well as some nuclear speckling (**Fig. 5D**). These trends are reflected in the N/C ratio measurements (**Fig. 5E**). Thus, the SMAD3 MH1 domain is not only sufficient but necessary as its disruption via the deletion of an α-helix inhibited SMAD3 signaling function and a loss of nuclear enrichment.

**Figure 5.**
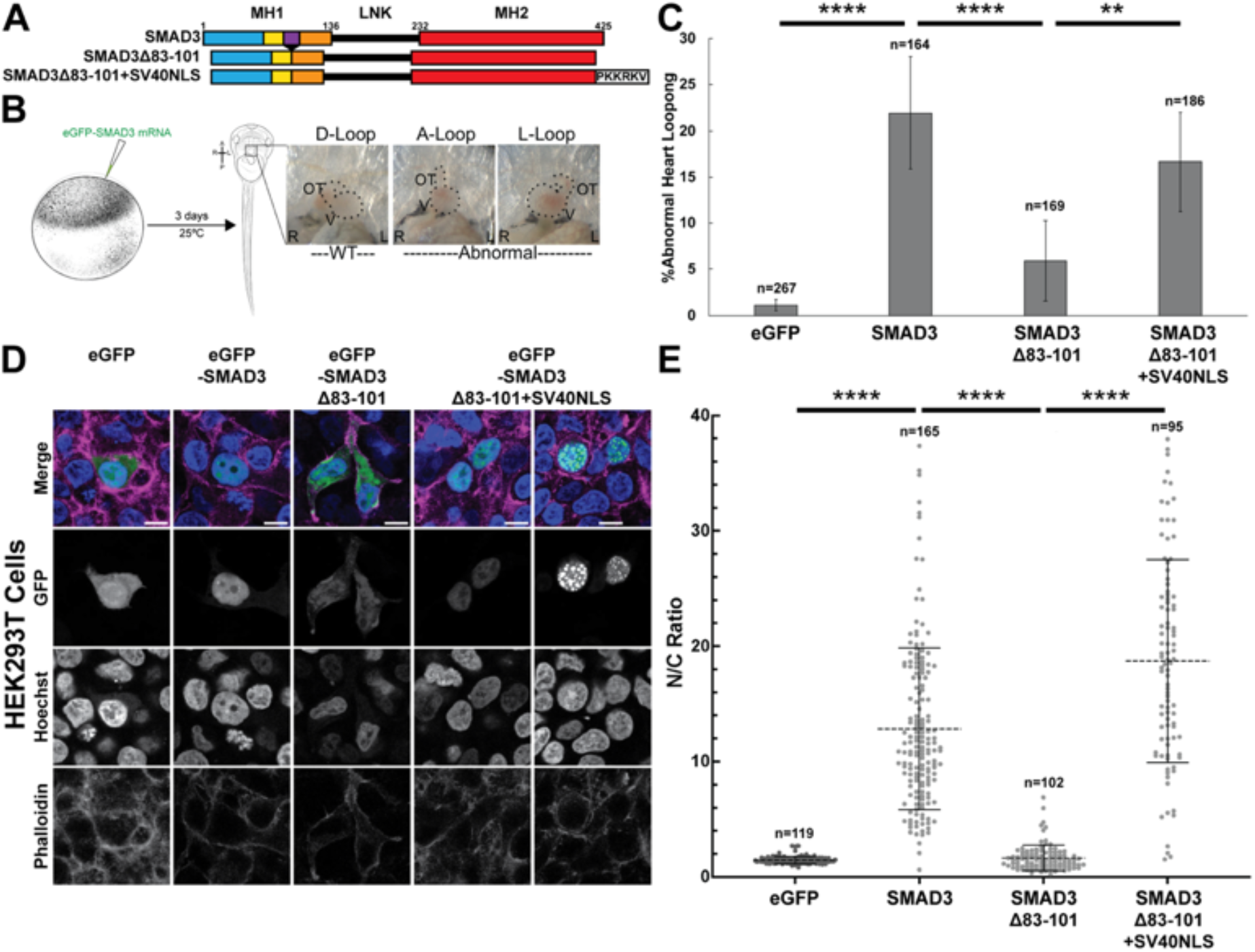
The identified SMAD NLS is functionally relevant in higher eukaryotes. **(A)** Schematic of eGFP-SMAD3 mRNAs. **(B)** Experimental workflow for functional testing of SMAD3 NLS in *X. tropicalis* embryos (left). (Right) Ventral views of *Xenopus* embryos with anterior to the top illustrate cardiac looping. Normal hearts loop to the right side of the animal (D-loop). Abnormally looped hearts can be un-looped (A-loop) or reversed (L-loop). *Xenopus* illustrations ©Natalya Zahn. **(C)** Percentage of embryos with abnormally-looped hearts. **** p<0.0001, *** p<0.0002, ** p<0.002, *p<0.02 (Fischer’s exact test). Error bars represent S.E.M. across 3 experiments (UIC 1.1±0.62%, eGFP-SMAD3 22.0±6.08%, eGFP-SMAD3Δ83-101 5.9±4.32%, eGFP-SMAD3Δ83-101+cNLS 16.7±5.37%). Total number of embryos from 3 biological replicates are reported above each error bar. **(D)** HEK293T cells expressing eGFP, eGFP-SMAD3, eGFP-SMAD3Δ83-101, or eGFP-SMAD3Δ83-101+cNLS. Cells were treated with Hoechst DNA stain and Phalloidin-A647 actin marker to label the nuclei and cell border, respectively. Scale bar=10 μm. Mean nuclear-to-cytoplasmic ratios of HEK293T cells expressing eGFP-SMAD3 mRNA. **** p<0.0001 (Unpaired T-test). Solid error bars represent standard deviation. Dashed line through scatterplot represents the mean (eGFP 1.5±0.03, SMAD3 14.7±12.33, Δ83-101 1.7±1.09, Δ83-101+cNLS 23.6±15.14) Experiments were done in triplicate and the total number of nuclei are labeled in the graph.

## DISCUSSION

In the current model of SMAD signaling, extracellular ligand binding to the cell-surface receptor leads to receptor heterodimerization. This dimerization leads to phosphorylation of an R-SMAD and subsequent formation of a heterotrimeric complex between R-SMADs and the Co-SMAD which accumulates in the nucleus activating target gene transcription^[2],[4]^. An important question is the status of SMAD nuclear transport without an external stimulus, which can be difficult to address in animal models. In addition, the behavior of individual SMADs without binding partners is difficult to address in multicellular animals. Understanding the basal nuclear transport mechanisms of an individual SMAD is a critical first step to understanding the activated context.

To address this question, we sought to develop a minimal model in which nuclear transport was conserved but lacked other components of TGF-β signaling, for which yeast is ideal. On the other hand, we cannot study the “activated” state when TGF-β ligands activate their respective receptors and initiate downstream signaling. Additionally, we lose potential interactions with other SMADs, other proteins, and possibly post-translational modifications that may not exist in yeast. Finally, there could be evolutionary differences in the structure of the NLSs/NESs across vertebrates and yeast or NTRs (for example at least one identified vertebrate NTR involved in import, Transportin2, does not have a yeast homolog). Nevertheless, our study found that SMAD2 and SMAD3 enrich in the nucleus while SMAD4, SMAD1 and SMAD5 are depleted. These results suggest that nuclear enrichment SMAD2/3 can occur without ligand-based activation. In the case of SMAD 1/5 and SMAD4, activation of the pathway must somehow suppress the tendency for these SMADs to otherwise remain in the cytoplasm. We were particularly struck by SMAD2/3’s nuclear enrichment in our yeast system and sought to investigate this further especially since nuclear entry in yeast does not require heterotrimeric complex formation as the prevailing model would propose.

To date, most studies identify importin-β, importin-7 and importin-8 as the SMAD2/3 nuclear importers^[7],[8],[13]^. However, a comprehensive survey of all known importins could have revealed additional players in SMAD2/3 nuclear enrichment. We decided to exclude Kap60 (importin-α) as there is no evidence that it acts independently of Kap95 (importin-β). Of the remaining 10 importins we tested, we found that only depletion of the yeast orthologs of importin-β, importin-7, and importin-8 (Kap95, Kap119, and Kap108) affected SMAD3 nuclear enrichment. This was reassuring and validated our approach in yeast. With this in hand, we next searched for the SMAD3 NLS.

Many studies have cited the KKLK sequence in the SMAD MH1 domain as the SMAD NLS^[8],[10],[11],[12],[13],[36],[37]^. There are two challenges with this putative NLS. First, at least one other study has shown that despite the necessity of this sequence, it is not sufficient to drive eGFP into the nucleus^[13]^. Second, this sequence is present in all R-SMADs and Co-SMADs, despite SMADs1, 4 and 5 being depleted from the nucleus. Our study showed that SMAD3(1-61), which contains this KKLK sequence, localizes exclusively to the cell periphery. We believe the cell-periphery localization may be due to the 2 α-helices that resemble amphipathic helices, which are known to interact with lipid bilayers^[38]^. We did, however, find that SMAD3(62-136) was sufficient to enrich eGFP in the nucleus. It is important to note that the MH1 domain is a minimally folding unit. Fragments of this domain expressed as in Figure 4 may not have retained the structures or functions they take when in the context of the full domain. Thus, it is likely that all the tested subdomains contribute to the MH1 domain’s nuclear enrichment capabilities. Functional testing in *X. tropicalis* and localization studies in mammalian cells showed that deleting an α-helix, SMAD3(83-101), led to a nuclear transport-specific loss of SMAD3 function. This all suggests that the necessary and sufficient SMAD3 NLS is more complex than the previously identified putative KKLK NLS.

NLSs are generally thought to be short peptides in IDRs which bind an NTR. Our results suggest that the entire MH1 domain, which is well structured, is required to confer robust nuclear import. This suggests that binding to the NTRs may require recognition of parts or all of the MH1 surface. There is precedent for such an idea. For example, an unusual heterodimer of importin-β and importin-7 engages Histone 1 much like a cradle^[39]^. Additionally, importin-8 binds eIF4e via electrostatic interactions in a folded domain^[40]^.

In our study we found that the full-length SMAD3 enriches in the nucleus. Interestingly, the MH1 domain fragment has a higher N/C ratio than full-length SMAD3. A potential explanation is that signaling-induced phosphorylation may cause a conformational change. This conformational change could make the NLS more available as in the case of the MH1 fragment. Alternatively, the conformational change could bury an NES^[11], [36]^. In the latter case, we might expect the linker or MH2 domains to contain an NES. However, when we tested these fragments, we did not find any evidence of an NES (Fig. 4).

The MH1 sequence we identified as the SMAD3 NLS is conserved across all of the SMADs, which would suggest that the other R-SMADs and Co-SMAD should also be similarly enriched in the nucleus, a challenge shared with the KKLK sequence from previous studies. One possibility is that differences in export may explain the different behaviors of the 3 classes of SMADs with regard to nuclear enrichment. Several studies have identified CRM1 as the primary exporter of SMAD1 and SMAD4^[10],[12],[16],[36]^. In addition, the SMAD4 NES (^142^DLSGLTLQ^149^) is in the linker region just following the MH1 domain and does not exist in any of the R-SMADs^[10],[16]^. Future studies could identify whether the observed nuclear depletion of the BMP SMADs is due to the use of particular exporters. A similar unbiased search for a BMP SMAD NES in our yeast system could also inform SMAD nuclear export mechanisms, given that we know it is ran-dependent in this system.

Our studies provide insight into nuclear transport as an important regulator for TGF-β signaling. Here we focus on the unstimulated distributions of SMADs. It is important to understand the basal behaviors of these signaling effectors in order to better investigate their behaviors in signaling conditions. In *S. cerevisiae*, SMAD 2 and 3 are able to enrich in the nucleus independently from any other SMADs and without the receptors required to phosphorylate and activate SMADs. Furthermore, BMP SMADs 1 and 5 and co-SMAD4 are depleted from the yeast nucleus. This knowledge of the basal nuclear distribution of SMADs allows us to ask better questions about what aspects of signaling control nuclear enrichment. In fact, our yeast system can be used to study basal nuclear transport mechanisms of other vertebrate signaling effectors including transcription factors offering exciting new avenues for the fields of nuclear transport, embryonic development, and disease.

## MATERIALS AND METHODS

### Contact for reagent and resource sharing

Additional information and requests for reagents may be directed to and fulfilled by the lead contact, Mustafa K. Khokha.See Supplementary File 1 for all strains, plasmids, and key resources.

#### Xenopus

*Xenopus tropicalis* are monitored, maintained, and cared for according to Yale University Institutional Animal Care and Use Committee (IACUC) protocols which are in line with ARRIVE guidelines. *Xenopus* lines have been housed and maintained in our aquatic animal facility for over 10 years. *In vitro* fertilizations were performed in accordance with established protocols^[41],[42],[43]^. All animal experimental protocols were approved by Yale’s IACUC and Animal Resource Center (YARC).

#### *S. cerevisiae* strains

All yeast strains used in this study are listed in Supplementary Table S1. Yeast strains were grown at 30ºC, room temperature (20-25ºC), or 37ºC in YPAD (1% bacto Yeast extract, 2% bactoPeptone, 2% glucose, and 0.25% Adenine sulfate). Transformation of yeast was carried out using standard protocols^[44]^.

#### Plasmids and mRNA

Primers and plasmids used in this study are listed in Supplementary Tables S2 and S3. Unless otherwise noted, all amino acid sequence numbering is based on the human SMAD3 sequence. Gibson Assembly (New England Biolabs, #E26115) was used to create a pRS406*ADH1* yeast expression vector containing an eGFP upstream of the multiple cloning site (MCS). Gibson Assembly (GA) was subsequently used to insert either the SV40 Large T-antigen cNLS (PKKRKV), full-length human SMADs, or SMAD truncations into the MCS to generate a yeast expression vector for N-terminally tagged SMADs. GA was also used to clone the N-terminally tagged SMADs out of the pRS406*ADH1* plasmid and insert it into the pCS107 vector for subsequent mammalian/*Xenopus* experiments. Primers for all GA were designed using NEBaseChanger with STOP codons added to the insert-vector junction of reverse primers. All GA reactions were done according to manufacturer instructions with molar ratios calculated using NEBioCalculator. Either GA or Q5 mutagenesis (NEB, #E0552S) was used to create SMAD3 truncations and mutate DNA-binding residues R74, Q76, and K81 to alanines from the pRS406*ADH1*or pCS107 plasmids. Q5 Insertion (Q5 Ins) was used to c-terminally tag pCS107_eGFP_hSMAD3Δ83-101 with the cNLS for vertebrate experiments. Primers for both Q5 reactions were designed using NEBase Changer and reactions were done according to manufacturer instructions. mRNAs were synthesized using the SP6 mMessage machine kit (Thermo Fisher Scientific, #AM1340) following manufacturer instructions.

#### Yeast Transformations

pRS406 expression plasmids containing the cNLS, full-length, or fragments of SMAD coding sequence under the control of the constitutively active *ADH1* promoter were transformed into the W303, Heh2-mCherry::NAT(BWCPL1314) strain, *mtr1-1::TRP1*, Heh2-mCherry::*KAN* (WHCPL16) strain, or one of the anchor away^[25]^ DTCPL1635, *Kap###-FRB::HIS3* strains (WHCPL1-12) at the *URA3* locus using the LiAC/ssDNA/PEG method^[44]^.

#### Yeast sub-cellular localization by microscopy

A yeast colony with genomic integration of the eGFP-tagged construct was cultured in 3mL YPAD overnight at either 30ºC (WT and AnchorAway;12-14hr) or RT (*mtr1-1*;14-16 hr) in YPAD, diluted to OD600 (0.5-0.25) in 12 mL YPAD, and left to grow to an OD600 (0.6-1.2) at either 30ºC (WT and AnchorAway; 4-6hr) or RT (*mtr1-1*; 6-8hr). 1mL of Anchor Away cultures were treated with either 1 μL of rapamycin stock solution (1 mg/mL dissolved in DMSO) or 1 μL carrier (DMSO) and incubated for 15 min at 30ºC prior to mounting on slides. By western blot, we detected some variability in eGFP-SMAD3 protein levels between Kap-FRB strains. However, there is no observable difference in protein levels between DMSO and Rapamycin treated samples within strains (Fig. S2) indicating that rapamycin treatment does not significantly affect protein amounts. To test permissive and restrictive temperatures for the *mtr1-1* strain, separate culture flasks were either incubated at RT or 37ºC for 1hr prior to mounting on slides. To prepare slides, 2.2 μL of the culture were re-suspended in CSM (Complete Supplement Mixture) Adenine + Glucose, mounted on a glass slide, covered with an 18X18 mm glass coverslip, and live imaged within 15 minutes of being removed from the incubator post-treatment. Slides were imaged using a DeltaVision wide-field microscope (GE Healthcare) with a CoolSnapHQ^2^ CCD camera. Images were deconvolved using the iterative algorithm of sofWoRx. 6.5.1 (Applied Precision, GE Healthcare).

#### *Xenopus* Left-Right Asymmetry Experiments

*Xenopus tropicalis* embryos were injected with a mix of 50 pg of various human SMAD3 mRNA constructs and mCherry mRNA (150-300pg) at the one-cell stage. Embryos were scored for cardiac looping via stereomicroscopy at Nieuwkoop and Faber (NF) stages 43-45.

#### Mammalian Cell Culture and Imaging

HEK293T cells (ATCC) were maintained and cultured with Dulbecco’s Modified Eagle Medium (DMEM) medium + 10% fetal bovine serum +1% penicillin and streptomycin in a T-25 flask. Cells were seeded in a 24-well plate on gelatin-coated coverslips (NeuVitro). Cells were transfected at 70-80% confluency using Lipofectamine 3000 (Thermo Fisher Scientific) following manufacturer protocols. 24-48 hr post-transfection, cells were fixed in 4% paraformaldehyde+PBS treated with Hoechst and Phalloidin-A647 and processed for immunofluorescence (Supplementary Table S4). Cells were tested for mycoplasma using MycoAlert Detection kit (Lonza). Cells were treated with Pro-Long Gold, mounted on slides, and cured overnight at 4ºC. Slides were imaged on a Leica Stellaris DIVE.

#### Quantification and Statistical Analysis

All fluorescent images were analyzed with Fiji. For quantification, the oval selection tool was used to draw a circular region of interest (ROI). Circles of equal size were used to create ROIs in the nucleus and in the cytoplasm of each cell. The Z-plane was kept the same for both ROIs. ROIs were then measured for mean fluorescence intensity, and nuclear-to-cytoplasmic (N/C) ratios were calculated. In the case of mammalian cell culture N/C ratios beyond 40 were excluded due to concerns of the potential effects of massive over-expression on cell health. Statistical significance was defined as <0.02 (*), <0.002 (**), <0.0002 (***) and <0.0001 (****). Unpaired two-tailed Students t-tests were used to determine statistical significance of N/C mean ratios in GraphPad Prism 10.0.1. Fischer’s exact tests were used to determine statistical significance of Abnormal HL rates in *X. tropicalis* embryos.

#### SMAD3 Structure Annotations

The SMAD3 full-length predicted structure was obtained from the AlphaFold Protein Structure Database (P84022)^[45],[46]^. SMAD3 crystal structures were viewed in the PDBe-KB and citations for original structures were obtained from the RSCB Protein Databank (RSCB.org, PDB IDs: 1OZJ, 1MJS, 1MK2, 5XOC, 5OD6, 5ODG, 6ZMN, 1U7F, 1MHD)^[26–32],[47–49]^. Structures were downloaded then annotated using PyMol V2.5.7.

## ACKNOWLEDGEMENTS

The authors thank the Yale CCMI core facilities for guidance using microscopes. Thanks to Elisa Rodriguez, Hue Tran, Maura Lane, and Michael Slocum for reagents and frog husbandry. Thank you to Dr. Yuh Min Chook for her feedback on this manuscript.

## AUTHOR CONTRIBUTIONS

[D.P.G.] Designed, conducted, visualized, analyzed, and validated experiments and wrote the original draft of and edited this manuscript. [M.C.E] Helped conduct, visualize, and analyze experiments in Figures 3 & 4. [C.P.L, M.K.K.] Supervised and guided project conception, experimental design, provided reagents, equipment, and funding, and edited this manuscript.

## DATA AND MATERIALS AVAILABILITY

Data, protocols, and reagents available upon request. Contact MKK for further information.

## SUPPLEMENTAL FIGURES

**Figure S1.**
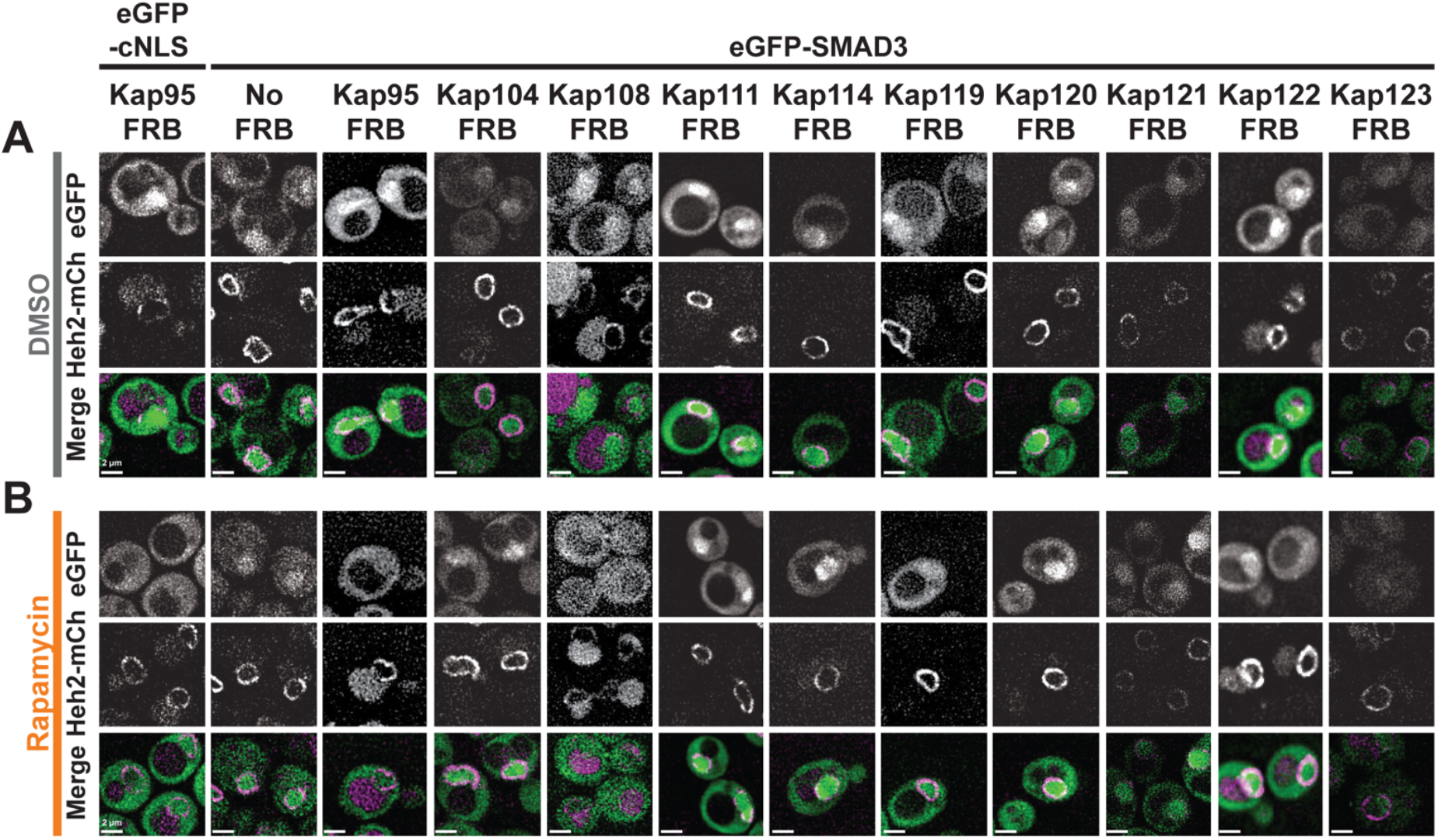
Images of eGFP-SMAD3 in NTR-FRB Anchor Away experiments. Anchor-away yeast strains co-expressing the nuclear envelope protein Heh2-mCherry and eGFP-SMAD3. eGFP-cNLS was also tested in the Kap95-FRB anchor-away as a positive control. **(A)** Cells imaged following a 15 min incubation with DMSO (gray). **(B)** Cells imaged following a 15 min incubation with Rapamycin (orange). Scale bar=2 μm.

**Figure S2.**
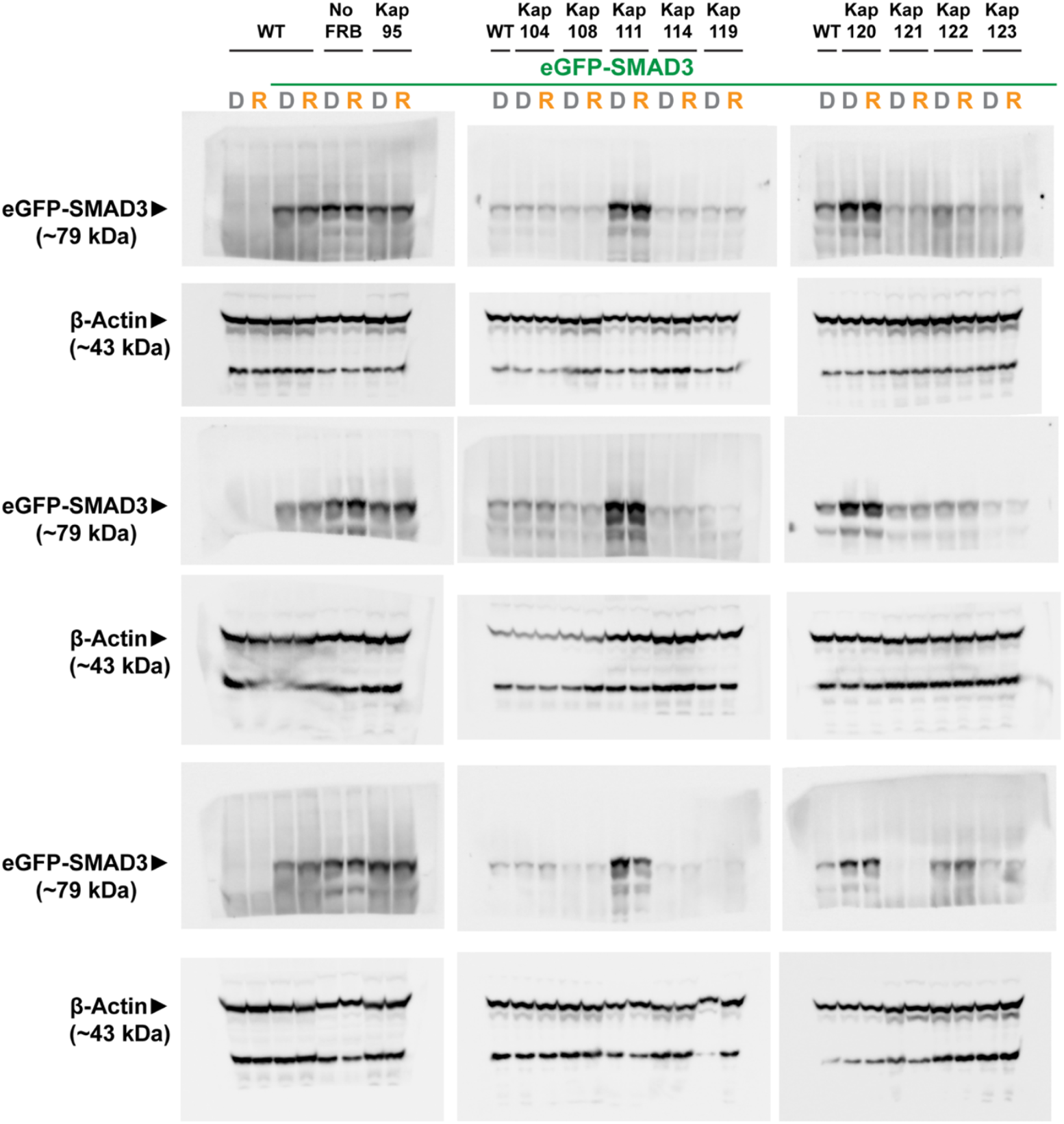
eGFP-SMAD3 expression levels in NTR Anchor-Away yeast strains. Protein was extracted from overnight cultures of wildtype Heh2-mCherry expressing yeast alone or co-expressing eGFP-SMAD3, and eGFP-SMAD3 expressing anchor-away yeast strains following a 15 min incubation with DMSO (D, gray) or rapamycin (R, orange). Whole protein extracts were run out on gels and transferred to a PVDF membrane. Membranes were cut at ~50kDa and probed for SMAD3 (~79 kDa for eGFP-SMAD3) or β-actin (~43 kDa). Each set of blots was from different overnight cultures. DMSO-treated wildtype Heh2-mCherry expressing yeast co-expressing eGFP-SMAD3 was included in the first lane for comparison across multiple blots.

## SUPPLEMENTAL METHODS

### Anchor Away Western Blots

1mL of overnight cultures were treated with either 1 μL of rapamycin stock solution (1 mg/mL dissolved in DMSO) or 1 μL carrier (DMSO) and incubated for 15 min at 30ºC prior to pelleting and re-suspended cultures in 1mM EDTA. Proteins were extracted by a 10 min incubation with ice-cold 2mM NaOH quenched with 50% trichloroacetic acid, wash with ice-cold acetone, 5 minute air-dry, and final denaturing in a 5% SDS Sample buffer with β-mercaptoethanol (40 uL/mL). Whole protein extracts were run out on gels and transferred to a PVDF membrane. Membranes were cut at ~50kDa and trimmed before treatment with primary antibodies to fit in the incubation container. The top half was treated with rabbit-α-SMAD3 (C67H9, 1:1000) primary and HRP-α-rabbit (1:10,000) secondary. The bottom half was treated with HRP-Rabbit-α-βActin (sc-47778, 1:10,000) primary antibody. Each set of blots was from different overnight cultures.

## SUPPLEMENTAL TABLES

**Supplementary Table 1.** Yeast Strains.

| Supplementary Table 1. Yeast Strains |  |  |  |
| --- | --- | --- | --- |
| Name | Genotype | Origin | Generation |
| W303a | MATa, *ade2-1 can1-100 his3-11,15 leu2-3,112 trp1-1 ura3-1* | EUROSCARF |  |
| W303α | MATα, *ade2-1 can1-100 his3-11,15 leu2-3,112 trp1-1 ura3-1* | EUROSCARF |  |
| HHY110 | W303, MAT *alpha tor1-1 fpr1::natMX6MX6 PMA1-2xFKBP12::TRP1* | Euroscarf (Haruki et al., 2008) |  |
| BWCPL1314 | W303, HEH2-mCherry::*natMX6* | (Thaller et al., 2019) |  |
| DPGCPL1 | BWCPL1314, eGFP::*URA3* | This study | Transformation of linearized pRS406ADH1_eGFP in BWCPL1314 |
| DPGCPL2 | BWCPL1314, eGFP-SV40NLS::*URA3* | This study | Transformation of linearized pRS406ADH1_eGFP_SV40NLS in BWCPL1314 |
| DPGCPL3 | BWCPL1314, eGFP-hSMAD1::*URA3* | This study | Transformation of linearized pRS406ADH1_eGFP_hSMAD1 in BWCPL1314 |
| DPGCPL4 | BWCPL1314, eGFP-hSMAD2::*URA3* | This study | Transformation of linearized pRS406ADH1_eGFP_hSMAD2 in BWCPL1314 |
| DPGCPL5 | BWCPL1314, eGFP-hSMAD3::*URA3* | This study | Transformation of linearized pRS406ADH1_eGFP_hSMAD3 in BWCPL1314 |
| DPGCPL6 | BWCPL1314, eGFP-hSMAD4::*URA3* | This study | Transformation of linearized pRS406ADH1_eGFP_hSMAD4 in BWCPL1314 |
| DPGCPL7 | BWCPL1314, eGFP-hSMAD5::*URA3* | This study | Transformation of linearized pRS406ADH1_eGFP_hSMAD5 in BWCPL1314 |
| DPGCPL8 | BWCPL1314, eGFP-hSMAD9::*URA3* | This study | Transformation of linearized pRS406ADH1_eGFP_hSMAD9 in BWCPL1314 |
| WHCPL16 | *mtr1-1::TRP1*, HEH2-mCherry::*KAN* | (Hwang et al., 2022) |  |
| DPGCPL9 | WHCPL16, eGFP::*URA3* | This study | Transformation of linearized pRS406ADH1_eGFP in WHCPL16 |
| DPGCPL10 | WHCPL16, eGFP-SV40NLS::*URA3* | This study | Transformation of linearized pRS406ADH1_eGFP_SV40NLS in WHCPL16 |
| DPGCPL11 | WHCPL16, eGFP-hSMAD1::*URA3* | This study | Transformation of linearized pRS406ADH1_eGFP_hSMAD1 in WHCPL16 |
| DPGCPL12 | WHCPL16, eGFP-hSMAD2::*URA3* | This study | Transformation of linearized pRS406ADH1_eGFP_hSMAD2 in WHCPL16 |
| DPGCPL13 | WHCPL16, eGFP-hSMAD3::*URA3* | This study | Transformation of linearized pRS406ADH1_eGFP_hSMAD3 in WHCPL16 |
| DPGCPL14 | WHCPL16, eGFP-hSMAD4::*URA3* | This study | Transformation of linearized pRS406ADH1_eGFP_hSMAD4 in WHCPL16 |
| DPGCPL15 | WHCPL16, eGFP-hSMAD5::*URA3* | This study | Transformation of linearized pRS406ADH1_eGFP_hSMAD5 in WHCPL16 |
| DTCPL1635 | HHY110, HEH2-mCherry::*KAN* | (Hwang et al., 2022) |  |
| WHCPL13 | DTCPL1635, *Kap95-FRB::HIS3*, | (Hwang et al., 2022) |  |
| WHCPL1 | DTCPL1635, *Kap104-FRB::HIS3*, | (Hwang et al., 2022) |  |
| WHCPL2 | DTCPL1635, *Kap108-FRB::HIS3* | (Hwang et al., 2022) |  |
| WHCPL3 | DTCPL1635, *Kap111-FRB::HIS3* | (Hwang et al., 2022) |  |
| WHCPL7 | DTCPL1635, *Kap114-FRB::HIS3* | (Hwang et al., 2022) |  |
| WHCPL10 | DTCPL1635, *Kap119-FRB::HIS3* | (Hwang et al., 2022) |  |
| WHCPL14 | DTCPL1635, *Kap120+FRB::HIS3* | (Hwang et al., 2022) |  |
| WHCPL15 | DTCPL1635, *Kap121+FRB::HIS3* | (Hwang et al., 2022) |  |
| WHCPL11 | DTCPL1635, *Kap122-FRB::HIS3* | (Hwang et al., 2022) |  |
| WHCPL12 | DTCPL1635, *Kap123-FRB::HIS3* | (Hwang et al., 2022) |  |
| DPGCPL16 | DTCPL1635, eGFP-hSMAD3::*URA3* | This study | Transformation of linearized pRS406ADH1_eGFP_hSMAD3 in DTCPL1635 |
| DPGCPL17 | WHCPL13, eGFP-hSMAD3::*URA3* | This study | Transformation of linearized pRS406ADH1_eGFP_hSMAD3 in WHCPL13 |
| DPGCPL18 | WHCPL1, eGFP-hSMAD3::*URA3* | This study | Transformation of linearized pRS406ADH1_eGFP_hSMAD3 in WHCPL1 |
| DPGCPL19 | WHCPL2, eGFP-hSMAD3:: *URA3* | This study | Transformation of linearized pRS406ADH1_eGFP_hSMAD3 in WHCPL2 |
| DPGCPL20 | WHCPL3, eGFP-hSMAD3:: *URA3* | This study | Transformation of linearized pRS406ADH1_eGFP_hSMAD3 in WHCPL3 |
| DPGCPL21 | WHCPL7, eGFP-hSMAD3:: *URA3* | This study | Transformation of linearized pRS406ADH1_eGFP_hSMAD3 in WHCPL7 |
| DPGCPL22 | WHCPL10, eGFP-hSMAD3:: *URA3* | This study | Transformation of linearized pRS406ADH1_eGFP_hSMAD3 in WHCPL10 |
| DPGCPL23 | WHCPL14, eGFP-hSMAD3:: *URA3* | This study | Transformation of linearized pRS406ADH1_eGFP_hSMAD3 in WHCPL14 |
| DPGCPL24 | WHCPL15, eGFP-hSMAD3:: *URA3* | This study | Transformation of linearized pRS406ADH1_eGFP_hSMAD3 in WHCPL15 |
| DPGCPL25 | WHCPL11, eGFP-hSMAD3:: *URA3* | This study | Transformation of linearized pRS406ADH1_eGFP_hSMAD3 in WHCPL11 |
| DPGCPL26 | WHCPL12, eGFP-hSMAD3:: *URA3* | This study | Transformation of linearized pRS406ADH1_eGFP_hSMAD3 in WHCPL12 |
| DPGCPL27 | BWCPL1314, eGFP-hSMAD3(1-136)::*URA3* | This study | Transformation of linearized pRS406ADH1_eGFP_hSMAD3(1-136) in BWCPL1314 |
| DPGCPL28 | BWCPL1314, eGFP-hSMAD3(137-231)::*URA3* | This study | Transformation of linearized pRS406ADH1_eGFP_hSMAD3(137-231) in BWCPL1314 |
| DPGCPL29 | BWCPL1314, eGFP-hSMAD3(232-425)::*URA3* | This study | Transformation of linearized pRS406ADH1_eGFP_hSMAD3(232-425) in BWCPL1314 |
| DPGCPL30 | BWCPL1314, eGFP-hSMAD3(1-61)::*URA3* | This study | Transformation of linearized pRS406ADH1_eGFP_hSMAD3(1-61) in BWCPL1314 |
| DPGCPL31 | BWCPL1314, eGFP-hSMAD3(62-136)::*URA3* | This study | Transformation of linearized pRS406ADH1_eGFP_hSMAD3(62-136) in BWCPL1314 |
| DPGCPL32 | BWCPL1314, eGFP-hSMAD3(62-136*)::*URA3* | This study | Transformation of linearized pRS406ADH1_eGFP_hSMAD3(62-136*) in BWCPL1314 |
| DPGCPL33 | BWCPL1314, eGFP-hSMAD3(62-101)::*URA3* | This study | Transformation of linearized pRS406ADH1_eGFP_hSMAD3(62-101) in BWCPL1314 |
| DPGCPL34 | BWCPL1314, eGFP-hSMAD3(62-82)::*URA3* | This study | Transformation of linearized pRS406ADH1_eGFP_hSMAD3(62-82) in BWCPL1314 |
| DPGCPL35 | BWCPL1314, eGFP-hSMAD3(83-101)::*URA3* | This study | Transformation of linearized pRS406ADH1_eGFP_hSMAD3(83-101) in BWCPL1314 |
| DPGCPL36 | BWCPL1314, eGFP-hSMAD3(102-136)::*URA3* | This study | Transformation of linearized pRS406ADH1_eGFP_hSMAD3(102-136) in BWCPL1314 |

**Supplementary Table 2.**
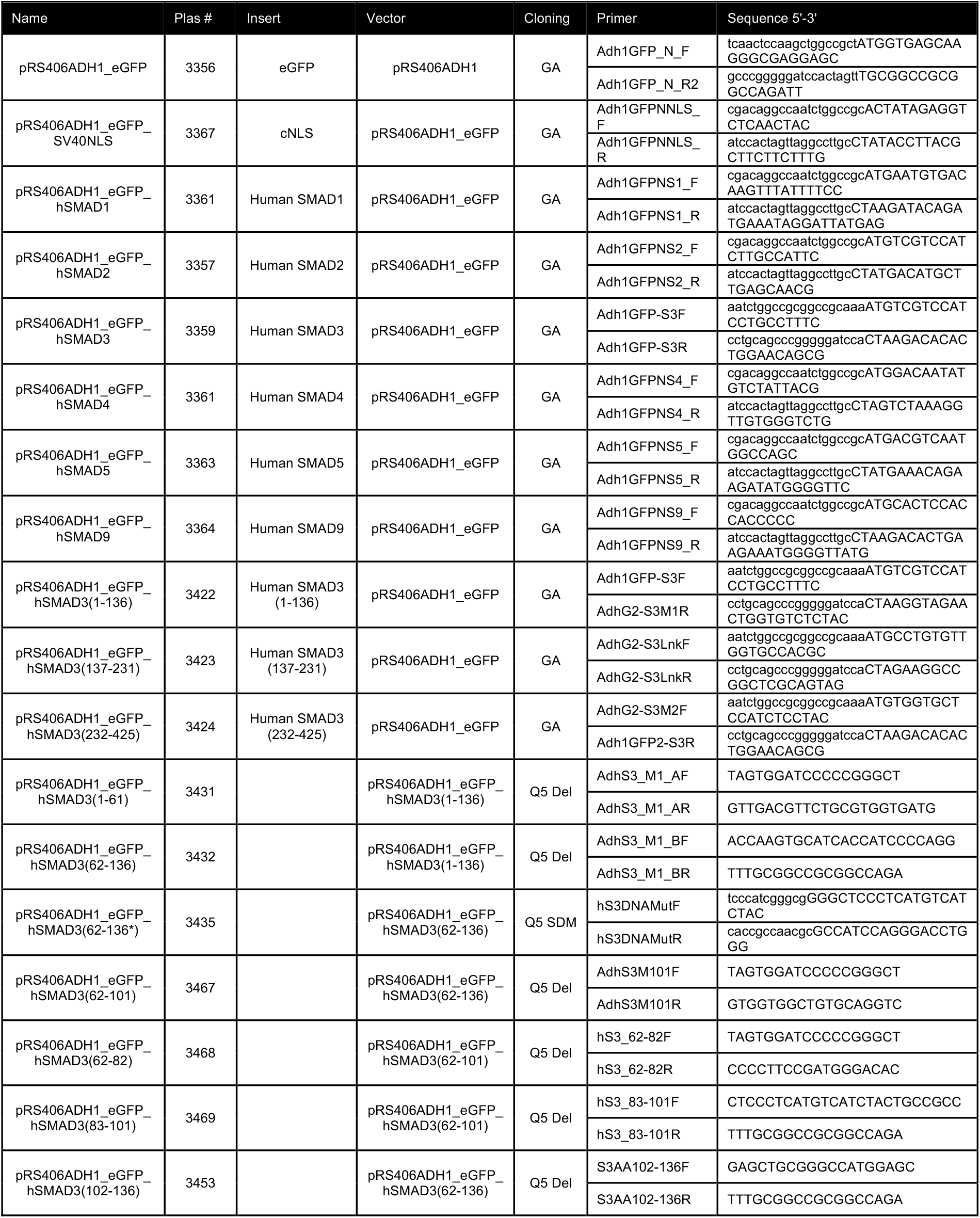
Plasmids for Yeast Experiments.

**Supplementary Table 3.**
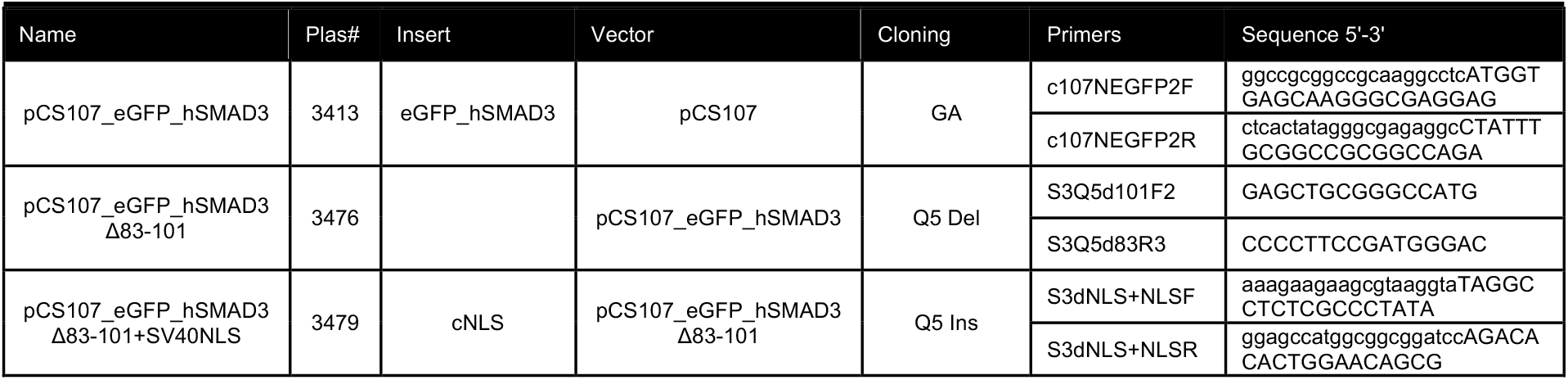
Plasmids for Vertebrate Experiments.

**Supplementary Table 4.**
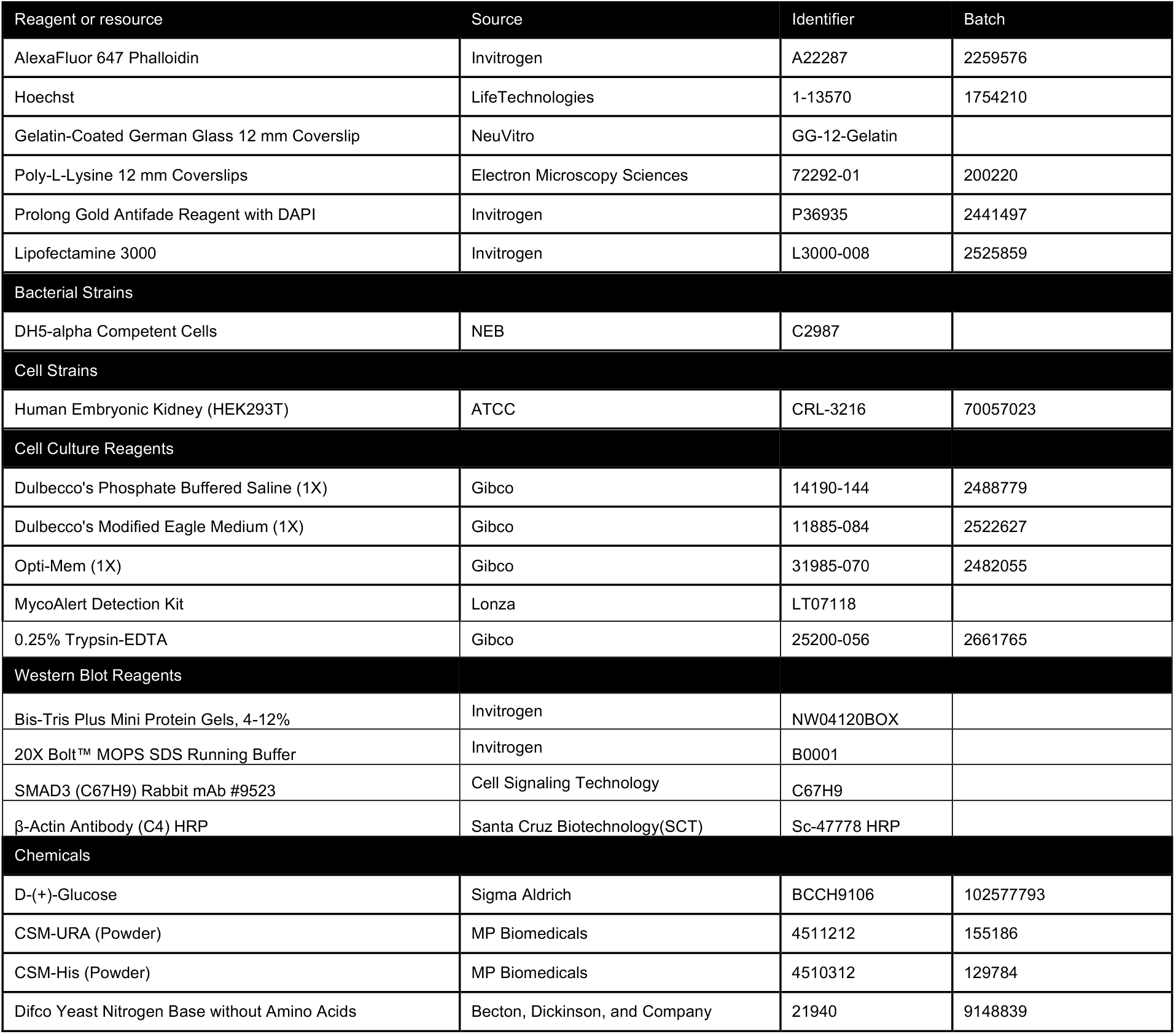
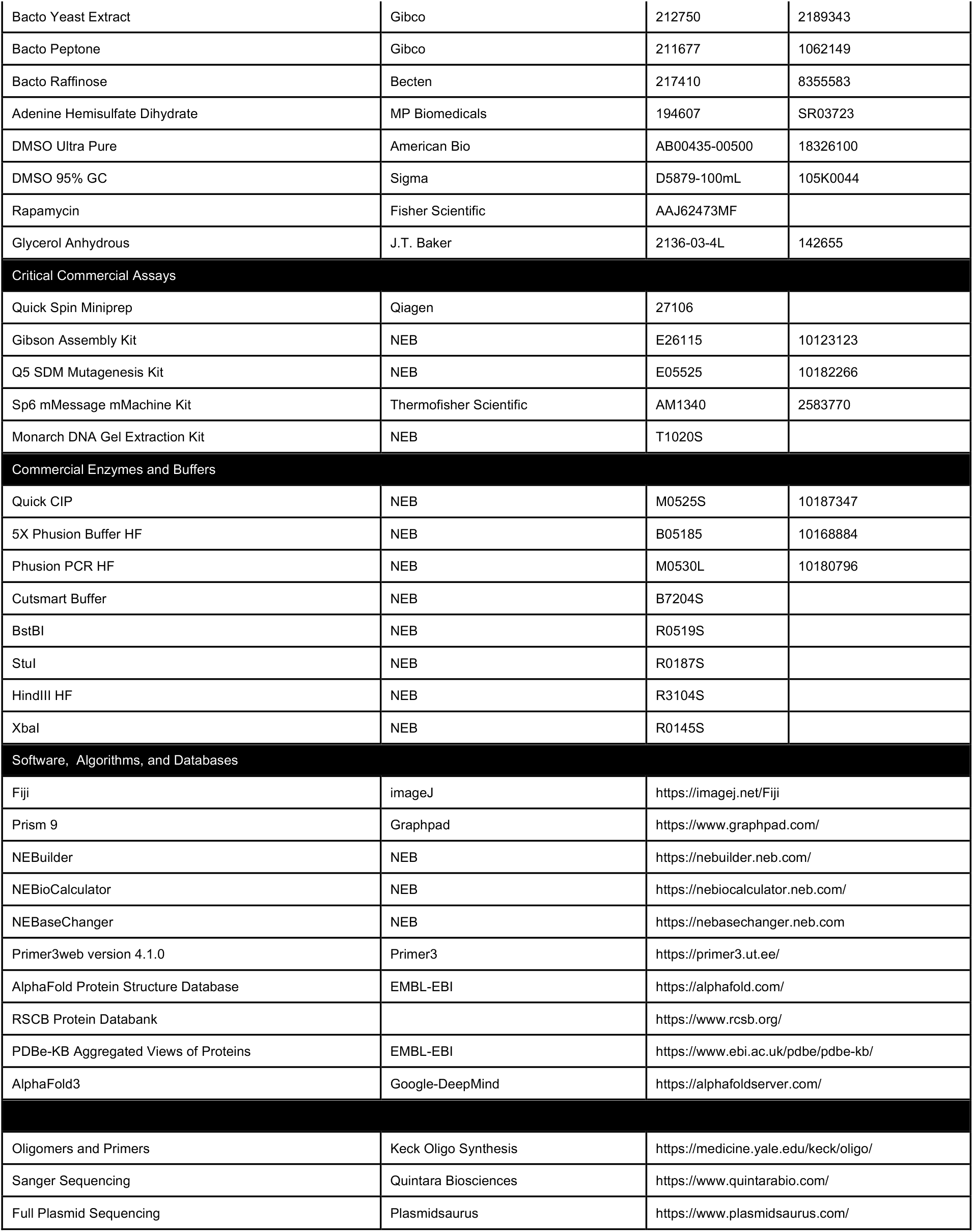
Key Resources.

## REFERENCES

[1] Hill, C. S. Nucleocytoplasmic shuttling of Smad proteins. Cell Res. 19, 36–46 (2009).

[2] Hill, C. S. Transcriptional control by the SMADs. Cold Spring Harb. Perspect. Biol. 8 (2016).

[3] Shi, Y. and Massagué, J. Mechanisms of TGF-β signaling from cell membrane to the nucleus. Cell. 113, 685–700 (2003).

[4] Massagué, J. and Sheppard, D. TGF-β signaling in health and disease. Cell. 186, 4007–4037 (2023).

[5] ing, C. E., Fung, H. Y. J. and Chook, Y. M. Karyopherin-mediated nucleocytoplasmic transport. Nat. Rev. Mol. Cell Biol. 23, 307–328 (2022).

[6] u, L., Yao, X., Chen, X., Lu, P., Zhang, B. and Ip, Y. T. Msk is required for nuclear import of TGF-β/BMP-activated Smads. J. Cell Biol. 178, 981–994 (2007).

[7] Kurisaki, A., Kose, S., Yoneda, Y., Heldin, C. H. and Moustakas, A. Transforming growth factor-β induces nuclear import of Smad3 in an importin-β1 and Ran-dependent manner. Mol. Biol. Cell. 12, 1079–1091 (2001).

[8] Xiao, Z., Liu, X. and Lodish, H. F. Acclerated Publication: Importin β mediates nuclear translocation of Smad 3. J. Biol. Chem. 275, 23425–23428 (2000).

[9] Kalderon, D., Richardson, W. D., Markham, A. F. and Smith, A. E. Sequence requirements for nuclear location of simian virus 40 large-T antigen. Nature. 311, 33–38 (1984).

[10] Pierreux, C. E., Nicolás, F. J. and Hill, C. S. Transforming Growth Factor β-Independent Shuttling of Smad4 between the Cytoplasm and Nucleus. Mol. Cell. Biol. 20, 9041–9054 (2000).

[11] Xiao, Z., Liu, X., Henis, Y. I. and Lodish, H. F. A distinct nuclear localization signal in the N terminus of Smad 3 determines its ligand-induced nuclear translocation. Proc. Natl. Acad. Sci. U. S. A. 97, 7853–7858 (2000).

[12] Xiao, Z., Latek, R. and Lodish, H. F. An extended bipartite nuclear localization signal in Smad4 is required for its nuclear import and transcriptional activity. Oncogene. 22, 1057–1069 (2003).

[13] Yao, X., Chen, X., Cottonham, C. and Xu, L. Preferential utilization of Imp7/8 in nuclear import of Smads. J. Biol. Chem. 283, 22867–22874 (2008).

[14] Borah, S., Thaller, D. J., Hakhverdyan, Z., Rodriguez, E. C., Isenhour, A. W., Rout, M. P., King, M. C. and Lusk, C. P. Heh2/Man1 may be an evolutionarily conserved sensor of NPC assembly state. Mol. Biol. Cell. 32, 1359–1373 (2021).

[15] Nakao, A., Imamura, T., Souchelnytskyi, S., Kawabata, M., Ishisaki, A., Oeda, E., Tamaki, K., Hanai, J., Heldin, C. H., Miyazono, K., et al. TGF-beta receptor-mediated signalling through Smad2, Smad3 and Smad4. EMBO J. 16, 5353–62 (1997).

[16] Watanabe, M., Masuyama, N., Fukuda, M. and Nishida, E. Regulation of intracellular dynamics of Smad4 by its leucine-rich nuclear export signal. EMBO Rep. 1, 176–182 (2000).

[17] Kadowaki, T., Zhao, Y. and Tartakoff, A. M. A conditional yeast mutant deficient in mRNA transport from nucleus to cytoplasm. Proc. Natl. Acad. Sci. U. S. A. 89, 2312–2316 (1992).

[18] Collignon, J., Varlet, I. and Robertson, E. J. Relationship between asymmetric nodal expression and the direction of embryonic turning. Nature. 381, 155–8 (1996).

[19] Hamada, H., Meno, C., Watanabe, D. and Saijoh, Y. Establishment of vertebrate left–right asymmetry. Nat. Rev. Genet. 3, 103–113 (2002).

[20] Kumar, A., Novoselov, V., Celeste, A. J., Wolfman, N. M., Ten Dijke, P. and Kuehn, M. R. Nodal signaling uses activin and transforming growth factor-β receptor-regulated Smads. J. Biol. Chem. 276, 656–661 (2001).

[21] Levin, M., Johnson, R. L., Sterna, C. D., Kuehn, M. and Tabin, C. A molecular pathway determining left-right asymmetry in chick embryogenesis. Cell. 82, 803–814 (1995).

[22] Lowe, L. A., Supp, D. M., Sampath, K., Yokoyama, T., Wright, C. V. E., Potter, S. S., Overbeek, P. and Kuehn, M. R. Conserved left–right asymmetry of nodal expression and alterations in murine situs inversus. Nature. 381, 158–161 (1996).

[23] Varlet, I. and Robertson, E. J. Left—right asymmetry in vertebrates. Curr. Opin. Genet. Dev. 7, 519–523 (1997).

[24] Haruki, H., Nishikawa, J. and Laemmli, U. K. The Anchor-Away Technique: Rapid, Conditional Establishment of Yeast Mutant Phenotypes. Mol. Cell. 31, 925–932 (2008).

[25] Hwang, W.Y., Kostiuk, V., González, D.P., Lusk, C.P., Khokha, M.K. Kap-β2/Transportin mediates β-catenin nuclear transport in Wnt signaling. eLife. 11, 1–25 (2022).

[26] Chacko, B. M., Qin, B. Y., Tiwari, A., Shi, G., Lam, S., Hayward, L. J., De Caestecker, M. and Lin, K. Structural basis of heteromeric smad protein assembly in TGF-beta signaling. Mol. Cell. 15, 813–23 (2004).

[27] Chai, J., Wu, J.-W., Yan, N., Massagué, J., Pavletich, N. P. and Shi, Y. Features of a Smad3 MH1-DNA complex. Roles of water and zinc in DNA binding. J. Biol. Chem. 278, 20327–31 (2003).

[28] Martin-Malpartida, P., Batet, M., Kaczmarska, Z., Freier, R., Gomes, T., Aragón, E., Zou, Y., Wang, Q., Xi, Q., Ruiz, L., et al. Structural basis for genome wide recognition of 5-bp GC motifs by SMAD transcription factors. Nat. Commun. 8, 2070 (2017).

[29] Miyazono, K.-I., Moriwaki, S., Ito, T., Kurisaki, A., Asashima, M. and Tanokura, M. Hydrophobic patches on SMAD2 and SMAD3 determine selective binding to cofactors. Sci. Signal. 11 (2018).

[30] Qin, B. Y., Lam, S. S., Correia, J. J. and Lin, K. Smad3 allostery links TGF-beta receptor kinase activation to transcriptional control. Genes Dev. 16, 1950–63 (2002).

[31] Ruiz, L., Kaczmarska, Z., Gomes, T., Aragon, E., Torner, C., Freier, R., Baginski, B., Martin-Malpartida, P., de Martin Garrido, N., Marquez, J. A., et al. Unveiling the dimer/monomer propensities of Smad MH1-DNA complexes. Comput. Struct. Biotechnol. J. 19, 632–646 (2021).

[32] Shi, Y., Wang, Y. F., Jayaraman, L., Yang, H., Massagué, J. and Pavletich, N. P. Crystal structure of a Smad MH1 domain bound to DNA: insights on DNA binding in TGF-beta signaling. Cell. 94, 585–94 (1998).

[33] Aragón, E., Wang, Q., Zou, Y., Morgani, S. M., Ruiz, L., Kaczmarska, Z., Su, J., Torner, C., Tian, L., Hu, J., et al. Structural basis for distinct roles of SMAD2 and SMAD3 in FOXH1 pioneer-directed TGF-β signaling. Genes Dev. 33, 1506–1524 (2019).

[34] BabuRajendran, N., Palasingam, P., Narasimhan, K., Sun, W., Prabhakar, S., Jauch, R. and Kolatkar, P. R. (2010). Structure of Smad1 MH1/DNA complex reveals distinctive rearrangements of BMP and TGF-β effectors. Nucleic Acids Res. 38, 3477–3488.

[35] Schweickert, A., Vick, P., Getwan, M., Weber, T., Schneider, I., Eberhardt, M., Beyer, T., Pachur, A. and Blum, M. The Nodal Inhibitor Coco Is a Critical Target of Leftward Flow in Xenopus. Curr. Biol. 20, 738–743 (2010).

[36] Xiao, Z., Watson, N., Rodriguez, C. and Lodish, H. F. Nucleocytoplasmic Shuttling of Smad1 Conferred by Its Nuclear Localization and Nuclear Export Signals. J. Biol. Chem. 276, 39404–39410 (2001).

[37] Xiao, Z., Brownawell, A. M., Macara, I. G. and Lodish, H. F. A Novel Nuclear Export Signal in Smad1 Is Essential for Its Signaling Activity. J. Biol. Chem. 278, 34245–34252 (2003a).

[38] Woolfson, D. N. Understanding a protein fold: The physics, chemistry, and biology of α-helical coiled coils. J. Biol. Chem. 299, 104579 (2023).

[39] Ivic, N., Potocnjak, M., Solis-Mezarino, V., Herzog, F., Bilokapic, S. and Halic, M. Fuzzy Interactions Form and Shape the Histone Transport Complex. Mol. Cell. 73, 1191–1203.e6 (2019).

[40] Volpon, L., Culjkovic-Kraljacic, B., Osborne, M. J., Ramteke, A., Sun, Q., Niesman, A., Chook, Y. M. and Borden, K. L. B. Importin 8 mediates m7G cap-sensitive nuclear import of the eukaryotic translation initiation factor eIF4E. Proc. Natl. Acad. Sci. U. S. A. 113, 5263–5268 (2016).

[41] Lane, M., Mis, E.K., Khokha, M.K. Obtaining Xenopus tropicalis Eggs. Cold Spring Harb Protoc. doi:10.1101/pdb.prot106344. (2022).

[42] Lane, M., Slocum, M., Khokha, M.K. Raising and Maintaining Xenopus tropicalis from Tadpole to Adult. Cold Spring Harb Protoc. doi:10.1101/pdb.prot106369. (2022).

[43] Lane, M., Khokha, M.K. Obtaining Xenopus tropicalis Embryos by In Vitro Fertilization. Cold Spring Harb Protoc. doi: 10.1101/pdb.prot106351. (2022).

[44] Amberg, D. C., Burke, D. J. and Strathern, J. N. Methods in Yeast Genetics: A Cold Spring Harbor Laboratory Course Manual. (Cold Spring Harbor Laboratory Press, 2005).

[45] Jumper, J., Evans, R., Pritzel, A., Green, T., Figurnov, M., Ronneberger, O., Tunyasuvunakool, K., Bates, R., Žídek, A., Potapenko, A., et al. Highly accurate protein structure prediction with AlphaFold. Nature 596, 583–589 (2021).

[46] Varadi, M., Anyango, S., Deshpande, M., Nair, S., Natassia, C., Yordanova, G., Yuan, D., Stroe, O., Wood, G., Laydon, A., et al. AlphaFold Protein Structure Database: Massively expanding the structural coverage of protein-sequence space with high-accuracy models. Nucleic Acids Res. 50, D439–D444 (2022).

[47] Berman, H. M., Westbrook, J., Feng, Z., Gilliland, G., Bhat, T. N., Weissig, H., Shindyalov, I. N. and Bourne, P. E. The Protein Data Bank. Nucleic Acids Res. 28, 235–42 (2000).

[48] Burley, S. K., Bhikadiya, C., Bi, C., Bittrich, S., Chao, H., Chen, L., Craig, P. A., Crichlow, G. V., Dalenberg, K., Duarte, J. M., et al. RCSB Protein Data Bank (RCSB.org): delivery of experimentally-determined PDB structures alongside one million computed structure models of proteins from artificial intelligence/machine learning. Nucleic Acids Res. 51, D488–D508 (2023).

[49] PDBe-KB consortium. PDBe-KB: collaboratively defining the biological context of structural data. Nucleic Acids Res. 50, D534–D542 (2022).

